# Phage-Antibiotic Synergy is Driven by a Unique Combination of Antibacterial Mechanism of Action and Stoichiometry

**DOI:** 10.1101/2020.02.27.967034

**Authors:** Carmen Gu Liu, Sabrina I. Green, Lorna Min, Justin R. Clark, Keiko C. Salazar, Austen L. Terwilliger, Heidi B. Kaplan, Barbara W. Trautner, Robert F. Ramig, Anthony W. Maresso

## Abstract

The continued rise in antibiotic resistance is precipitating a medical crisis. Bacteriophage (phage) has been hailed as one possible therapeutic option to augment the efficacy of antibiotics. However, only a handful of studies have addressed the synergistic relationship between phage and antibiotics. Here, we report a comprehensive analysis of phage-antibiotic interaction that evaluates synergism, additivism, and antagonism for all classes of antibiotics across clinically achievable stoichiometries. We combined an optically-based real-time microtiter plate readout with a matrix-like heatmap of treatment potencies to measure phage and antibiotic synergy (PAS), a process we term synography. Phage-antibiotic synography was performed against a pandemic drug-resistant clonal group of *E. coli* (ExPEC) with antibiotic levels blanketing the minimum inhibitor concentration (MIC) across seven orders of viral titers. Our results suggest that, under certain conditions, phages provide an adjuvating effect by lowering the MIC for drug-resistant strains. Furthermore, synergistic and antagonistic interactions are highly dependent on the mechanism of bacterial inhibition by the class of antibiotic paired to the phage, and when synergism is observed, it suppresses the emergence of resistant cells. Host conditions that simulate the infection environment, including serum and urine, suppress PAS in a bacterial growth-dependent manner. Lastly, phage burst size seems to be a significant driver of synergism. Collectively, this data suggests lytic phages can resuscitate an ineffective antibiotic for previously resistant bacteria, while also synergize with antibiotics in a class-dependent manner, processes that may be dampened by lower bacterial growth rates found in host environments.

**Significance Statement:** Bacteriophage (phage) therapy is a promising approach to combat the rise of multi-drug resistant bacteria. Currently, the preferred clinical modality is to pair phage with an antibiotic, a practice thought to improve efficacy. However, antagonism between phage and antibiotics has been reported, the choice of phage and antibiotic is not often empirically determined, and the effect of the host factors on the effectiveness is unknown. Here, we interrogate phage-antibiotic interactions across antibiotics with different mechanisms of action. Our results suggest that phage can lower the working MIC for bacterial strains already resistant to the antibiotic, is dependent on the antibiotic class and stoichiometry of the pairing, and is dramatically influenced by the host microenvironment.

## Introduction

A major public health crisis is the alarming increase of infections caused by antibiotic-resistant bacteria, which are responsible for approximately 2.8 million infections in the U.S. alone (1). In 2014, the World Health Organization (WHO) reported that a post-antibiotic era, in which antibiotics are largely ineffective, is a possible fate for the 21^st^ century (2). The postantibiotic era is described as a period where antibiotics fail to target multidrug-resistant bacteria. Five years later in 2019, the Centers for Disease Control and Prevention (CDC) in its Antibiotic Resistance Threat Report claimed that the post-antibiotic era had already arrived (1). In a report released by a U.K. Commission, it was concluded that 10 million deaths a year, at a cost of around 3 trillion dollars, will occur due to drug-resistant infections by the year 2050 (3). Furthermore, the exchange of genetic elements that confer resistance is common amongst members of the human microbiome, resistance has developed against every major chemical class of antibiotics, and the overuse of antibiotics in patients and agriculture selects for such strains.

Of great concern is the pandemic clonal group called <u>ex</u>traintestinal <u>p</u>athogenic *<u>E</u>. <u>c</u>oli* (ExPEC) with the sequence type 131 (ST131) (4). This group demonstrates a highly virulent phenotype and is a prominent cause of urinary tract, peritoneal, bloodstream, and neonatal meningitis infections, while also being resistant to fluoroquinolone and **β**-lactam antibiotics (5, 6). In addition, ExPEC is a highly versatile pathogen comprised of many additional circulating sequence types possessing a plethora of virulence and resistance genes (4).

To address this growing problem, several alternatives to traditional chemical antibiotics have been explored: antibody therapy, antimicrobial peptides, probiotics, metal chelation and even incentives to expedite the drug-approval process and stimulate new antibiotic development (7). Although promising, all of these approaches are limited by the fact that the antimicrobial agent cannot change or adapt in real-time. That is, should resistance arise, the agent has little ability to become effective again, especially considering the significant investment in time and dollars needed to bring new drugs to the market. The ability of bacteria to mutate quickly and evolve around such approaches are both their greatest asset and a biological reality that disincentivizes investment in antibiotic-making business. In contrast, bacteriophages, viruses that infect and kill bacteria, are as equally evolvable and adaptable as bacteria, in addition to being the most numerous replicating entity on Earth (estimated to be around 10^31^ total particles)(8).

The adaptability and sheer number of phages imply that they are the largest repository of antibacterial information available to modern medicine. Furthermore, phages have been used to treat bacterial infections for decades in Eastern Europe, and recent compassionate care cases in the U.S. and U.K. has demonstrated clinical success (9, 10). Phages are also specific for a given bacterial species (even strain), meaning off-target killing of “good” bacteria in our microbiome can be minimized. Phages also amplify at the site of the infection, and therefore self-dose, clearing when no longer needed due to excretion or breakdown in the host (11). Finally, phages are generally regarded as safe by the food and drug administration (FDA), with millions of phage particles ingested every day in our food and water (12). These characteristics allow phages to become a potentially better alternative than chemical compounds for therapy, which, if developed through the normal pharma pipeline, may take ten years and a billion dollars to bring to market.

One of the more attractive and feasible use of phages is to combine them with clinically-used antibiotics, a sort of one-two punch on pathogenic bacteria (13). The combined use of phage and antibiotics may result in a number of outcomes. The two agents may act additively, that is, the sum of their individual effects is equal to their combinatorial efficacy. They may also act synergistically; their total efficacy is much greater than each individual action. A third result is no effect, owing to the lack of action of each individual agent. Finally, there may be antagonism whereby the molecular action of one of the agents somehow interferes with the action of the other. In reported cases of phage therapy in the U.S., the choice of antibiotic was often made on the bases of antibiogram data and the medical condition of the patients (14). However, the recent awareness of antagonism has propelled some *in vitro* assessment of phage-antibiotic action prior to treatment to select for synergistic combinations, a personalized approach that has led to satisfactory therapeutic outcomes (15, 16). Several phage-antibiotic combinations have been investigated *in vitro* and *in vivo* in multiple bacterial species (17, 18) but there have been mixed results with combinatorial treatment (13, 19, 20). For instance, quinolones can be synergistic with phages against *P. aeruginosa* in one study while antagonistic in another (21, 22). Sometimes, there are even two types of interactions found with the same antibiotic when they are combined with phages (23). Moreover, phage-antibiotic synergy (PAS) is usually studied with only one or two concentrations of the antimicrobials, which are wholly insufficient in predicting combinatorial concentrations that are efficacious during treatment. For these reasons, we assessed the effect of lytic phage on bacterial killing as a function of the presence of a resistance gene, the mechanism of action of the antibiotic, the likelihood of resistance, and the influence of host environments on effectiveness of PAS. We did these studies with a characterized myovirus (ΦHP3) that targets a pandemic clonal group of highly virulent ExPEC (strain JJ2528). Our investigation suggests that for phage-antibiotic combination therapy, clinicians should 1) pair antibiotics with phages whose production machinery within the bacterial cell does not rely on the bacterial process that are inhibited by the very antibiotics they wish to use, 2) consider the stoichiometry of the interactions, and 3) take into account how the host environment may affect treatment efficacy.

## Results

### Formation of a comprehensive antibiotic-phage synergy system – the synogram

To understand the range of possible outcomes on a bacterial growth when exposed to both phage and antibiotic, we assessed bacterial growth when exposed to concentrations of antibiotics that blankets the MIC across multiple orders of magnitude of phage titer over time in an optically-based microtiter plate assay system. The primary phage used throughout the investigation, ΦHP3, is a highly effective killer of the ST131 ExPEC clinical isolates and is commonly used as a prototype in the laboratory (24). Since an inoculum effect was observed with this isolate in the presence of different antibiotics (Fig. S2), bacteria were seeded at high inoculum to allow possible interaction between phage and antibiotic to occur at sub-inhibitory conditions. The optical density of the culture was then monitored at body temperature for twenty-four hours. The absorbance was read as a stand-alone parameter and converted to a heat map that represents the percentage of reduction of the bacterial population, what we refer to as a synogram (Fig. 1A and Fig. S3 – *raw data*).

**Figure 1.**
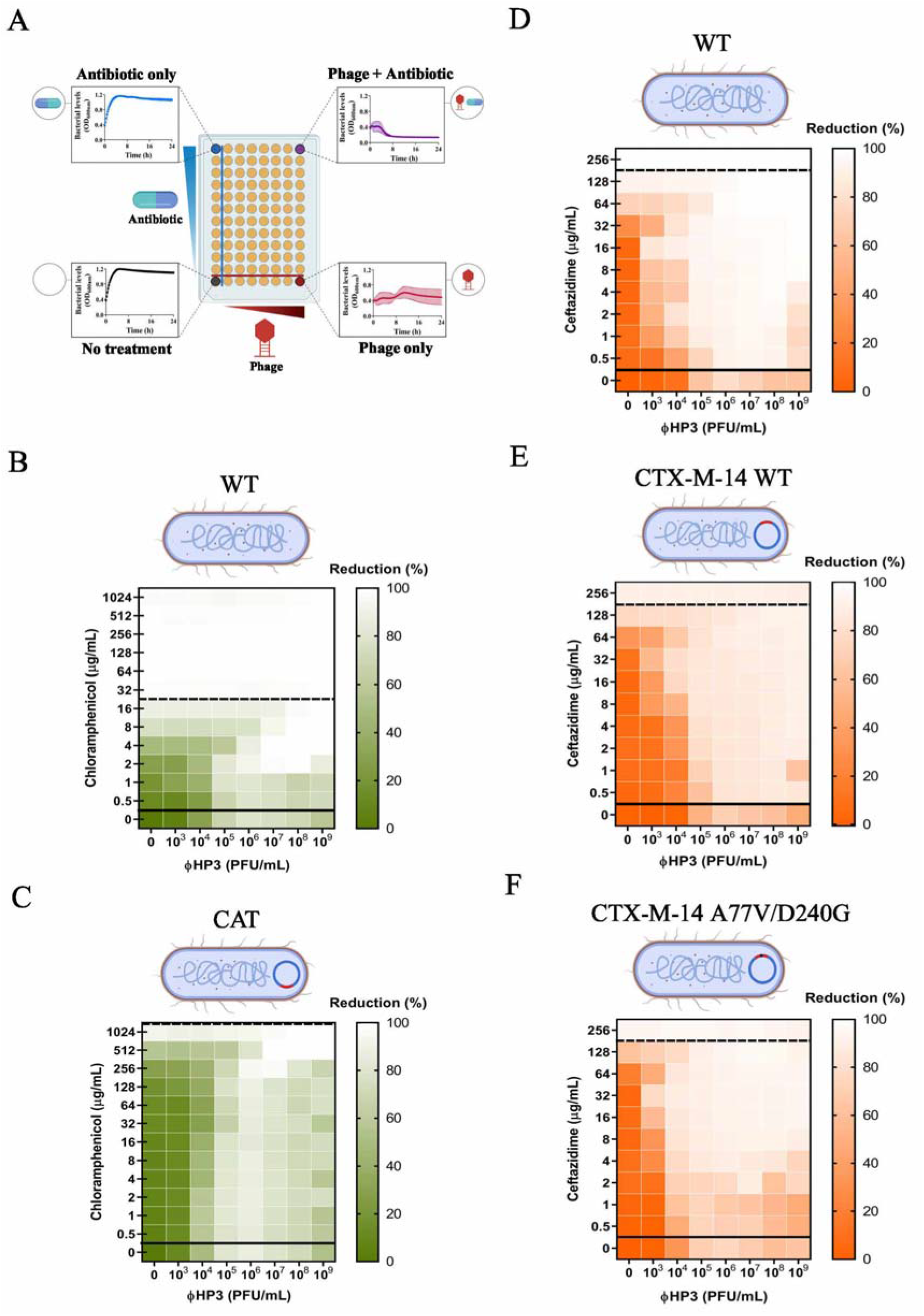
The effect of the bacterial resistance on phage-antibiotic synergy. A 100-fold diluted sub-culture of JJ2528 was incubated for 4 hours, centrifuged, washed, adjusted to O.D._600nm_ of 1, and inoculated onto a 96-well plate to which different treatments had been added to each well: phage alone (ΦHP3), antibiotic alone, phage-antibiotic combined, and untreated control. The OD_600nm_ was measured every 15 minutes for a total of 24 hours at 37°C with shaking. (**A**) Synogram showing different treatments. The effect of antibiotic-resistance on the gene and allele levels were shown as follow: chloramphenicol-HP3 combined treatment on (**B**) chloramphenicol sensitive JJ2528 and (**C**) chloramphenicol resistant JJ2528; ceftazidime-ΦHP3 combined treatment on (**D**) wild type JJ2528, (**E**) JJ2528 CTX-M-14 wild type, and (**F**) JJ2528 CTX-M-14 A77V/D240G. Synograms represent the mean reduction percentage of each treatment from three biological replicates: *Reduction* % = [(*OD_growth control_*) – (*OD_treatment_*)]*x*100. The region above the dashed line indicates antibiotic-mediated killing with highly effective doses; the region between the solid and dashed lines represents the interacting region of the phage and antibiotic, and the region below the solid line indicates phage-mediated killing with ineffective antibiotic concentrations.

Throughout the study, we found that synograms seemed to follow a pattern unique to the antibiotic being tested and can be generally divided into three sections: i) an antibiotic-dominated-killing region, usually the upper division of the synograms, where antibiotics are effective and thus the killing pattern of the combination therapy closely tracks the antibiotic-only treated cells (Figure 1A, *upper left hand corner*); ii) an interacting region, the middle section of the synograms, where the effect is a combination of both phage and antibiotic (additive, synergistic, or antagonistic – Fig. 1A, *middle to upper right hand corner*); and lastly, iii) a phage-dominated killing region, the lower segment of the synograms, where antibiotics are ineffective and the killing activity is influenced more by the phage. Representing the data as a synogram achieves two main objectives that would not be realized by scanning the plethora of information generated from such data sets. First, it allows for a convenient colorimetric visualization of the effectiveness of phage-antibiotic interactions as their concentrations change relative to each other. By simply looking for points of low intensity, it allows one to determine the optimal concentration of each agent for maximal killing. Second, the synograms allow for an easy comparison of the global effectiveness of any antibiotic-phage combination across multiple agents and conditions. This includes an assessment of different classes of antibiotics, different phage sequence types, and under different conditions, including those designed to simulate the host. One useful quantitative parameter when determining a phage-antibiotic concentration to use is the pairing of the two that reduces bacterial density by 90% or greater, a value that is denoted as the Synogram10 (Sn_10_). This way, multiples of the Sn_10_ (2X, 5X, etc.) can be used as a practical parameter when defining the concentration of each for an assay, experiment, or treatment.

### The effect of bacterial antibiotic resistance on combined phage-antibiotic efficacy

Using the synogram as a proxy for combinatorial efficacy, we first asked how PAS may change when the only difference in the system is the absence or presence of an antibiotic-resistance gene. For this, we assessed the phage-antibiotic killing dynamics of ExPEC strain JJ2528 lacking or containing the gene encoding the enzyme chloramphenicol acetyltransferase (CAT). CAT functions to transfer an acetyl group from coenzyme A to chloramphenicol, a modification that prevents the antibiotic from binding to the bacterial ribosome, prohibiting the inhibition of protein synthesis. We first examined the effect of each agent alone and in combination on wild-type JJ2528 that lacks the *cat* gene. The synogram of wild type JJ2528 shows almost complete reduction (>95%) when bacterial cells were treated with ≥ 32 μg/mL of chloramphenicol, a region that is predominantly affected by the action of the antibiotic (Fig. 1B, above dashed line). In the ΦHP3-alone treated cells, there was a progressive reduction from 10^3^-10^6^ PFU/mL. At higher phage titers, bacterial cells seemed to increase as phage titer increased (Fig. 1B, below solid line). This equates to a second growth of bacterial cells, likely resistance to the phage (examined below), a phenomenon that was observed throughout the study. Under combinatorial treatment, there is a nearly complete reduction of wild-type JJ2528 with concentrations as low as 1 μg/mL of chloramphenicol, suggesting that the addition of phage reduced the effective MIC of chloramphenicol 32-fold. The antibiotic, however, did not seem to appreciably reduce the phage titer needed for effective killing.

On the other hand, ExPEC harboring the enzyme CAT (JJ2528-CAT) is highly resistant to chloramphenicol; it takes >32-fold more (1024 μg/mL) chloramphenicol to achieve a similar percent of reduction as the wild-type JJ2528 that lacks the enzyme (Fig. 1C, compare the leftmost column to the same column in Fig. 1B). Once again, despite the presence of the CAT enzyme, combinatorial phage-antibiotic treatment results in a similarly high level of reduction of bacterial density with low chloramphenicol dose, with a general downward trend in reduction observed to the lowest dose of chloramphenicol (0.5 μg/mL). As is true for the CAT-minus ExPEC, phage seems to enhance chloramphenicol-based killing of the bacteria with little to no stimulation of the antibiotic on phage-based killing. In this regard, and as analyzed via the use of interaction plots (which allows one to determine if the effect is additive, synergistic, or antagonistic – see Fig. 3G), this type of phage-antibiotic killing can be described as an additive effect for both JJ2528 harboring either a chloramphenicol resistance gene or not. Interestingly, the interacting region for this synogram is highly dominated by phage killing with subtle variations in some wells. There were several combinations of lower titer of phage and higher doses of chloramphenicol in both wild type JJ2528 and JJ2528-CAT that yielded subtle antagonistic interactions (Fig. 1B, 1C and 3G). Interactive regions of the synograms were analyzed by determining the area under the curve (AUC) and plotted as violin plots (Fig. S1). These plots allow the visualization of the differences between synograms as a whole and also they take into account how bacterial populations, subjected to different treatments, change over time. In general, the addition of a resistance gene shifted more of the combined treatment matrix to the right (a lower % reduction – Fig. S1A, green).

The acquisition of a gene (like above) that encodes an enzyme or other protein that inactivates, blocks or pumps out the antibiotic is one mode by which bacteria become resistant to antibiotics. Another mode, however, is that the gene acquires mutations that enhance the corresponding enzyme’s catalytic activity. To determine the effect of phage-antibiotic combinatorial treatment on bacteria harboring such changes, we introduced genes encoding the **β**-lactamases CTX-M-14 WT and CTX-M-14 A77V/D240G into JJ2528. CTX is an enzyme that hydrolyzes the **β**-lactam ceftazidime; the presence of the double mutations (A77V/D240G) allows the bacteria to hydrolyze ceftazidime more efficiently than the wild type enzyme (25). Mutant versions of CTX-M are correlated with clinically high rates of resistance and serve here as both a relevant and controlled model to determine the effect phage may have on treatment (26–28). In contrast to the previous introduction of CAT, which produced sharply defined resistance, the introduction of these beta-lactamases yielded a subtle increase of resistance against ceftazidime, with an increase in the MIC of about 2-fold (Fig. 1D-F). In general, the ceftazidime synograms showed a single large interacting region. Many phage-antibiotic combinations efficiently killed planktonic cells, unlike the situation with chloramphenicol. In all three, a high degree of reduction was observed when phage was combined even with low antibiotic concentrations (0.5 μg/mL for JJ2528 WT, 0.5 μg/mL for JJ2528 CTX-M-14 WT, and 2 μg/mL for JJ2528 CTX-M-WT A77V/D240G), as opposed to ceftazidime-alone treated cells (128-256 μg/mL for all three). Consistent with the synogram, there was more killing of JJ2528 WT (Fig. 1D) than of JJ2528 CTX-M-14 WT (Fig. 1E), while JJ2528 CTX-M-14 A77V/D240G showed the least reduction of bacteria at combinatorial concentrations (Fig. 1F, Fig. S1A, orange). However, the addition of the **β**-lactamase gene to wild-type cells had much less of a total effect on bacterial levels than did the addition of chloramphenicol acetyltransferase (compare Fig. 1B and 1C to 1D and 1E). Unlike that observed for chloramphenicol, ceftazidime and ΦHP3 were synergistic over most of the concentrations of both agents, with as little as 1,000 PFU/mL reducing the MIC of the ceftazidime by 32-fold (Fig. 1D-F and Fig. 3F). This raises interesting questions as to how the specific mechanism of action of an antibiotic may also affect its ability to synergize with phage.

### The relationship between antibiotic mechanism of action and PAS

The observation that two different classes of antibiotics (protein synthesis versus cell wall synthesis inhibitors) showed dramatically different interactions with the same phage over the same concentration of both agents suggested that the outcome of phage-antibiotic interactions might vary in a consistent manner by antibiotic class (or mechanism of action). To test this hypothesis, we determined the effect on JJ2528 when ΦHP3 is combined with representatives of four other classes of antibiotics, including trimethoprim (folic acid synthesis inhibitor), colistin (cell membrane disrupter), ciprofloxacin (DNA topoisomerases inhibitor), and kanamycin (protein synthesis inhibitor-30S subunit). Wild-type JJ2528 treated with ΦHP3 and the folic acid synthesis inhibitor trimethoprim produced a synogram dominated by phage killing (Fig. 2A). In fact, JJ2528 was not efficiently killed even on high trimethoprim concentrations (for example, 256 μg/mL). However, a reduction was achieved with either phage-alone treated cells (≥10^4^ PFU/mL) or phage-antibiotic treated cells (Fig. 2A). Of note, the combined highest dose (256 μg/mL of trimethoprim and 10^7-9^ PFU/mL of ΦHP3) seemed to be able to clear more bacteria than either treatment alone (Fig. 2A); however, when this combination was analyzed further by examining the growth curve and corresponding interaction plots, these differences were not statistically significant (Fig. 3B). This type of pairing can thus be classified as no effect (Fig. 3B, interaction plots).

**Figure 2.**
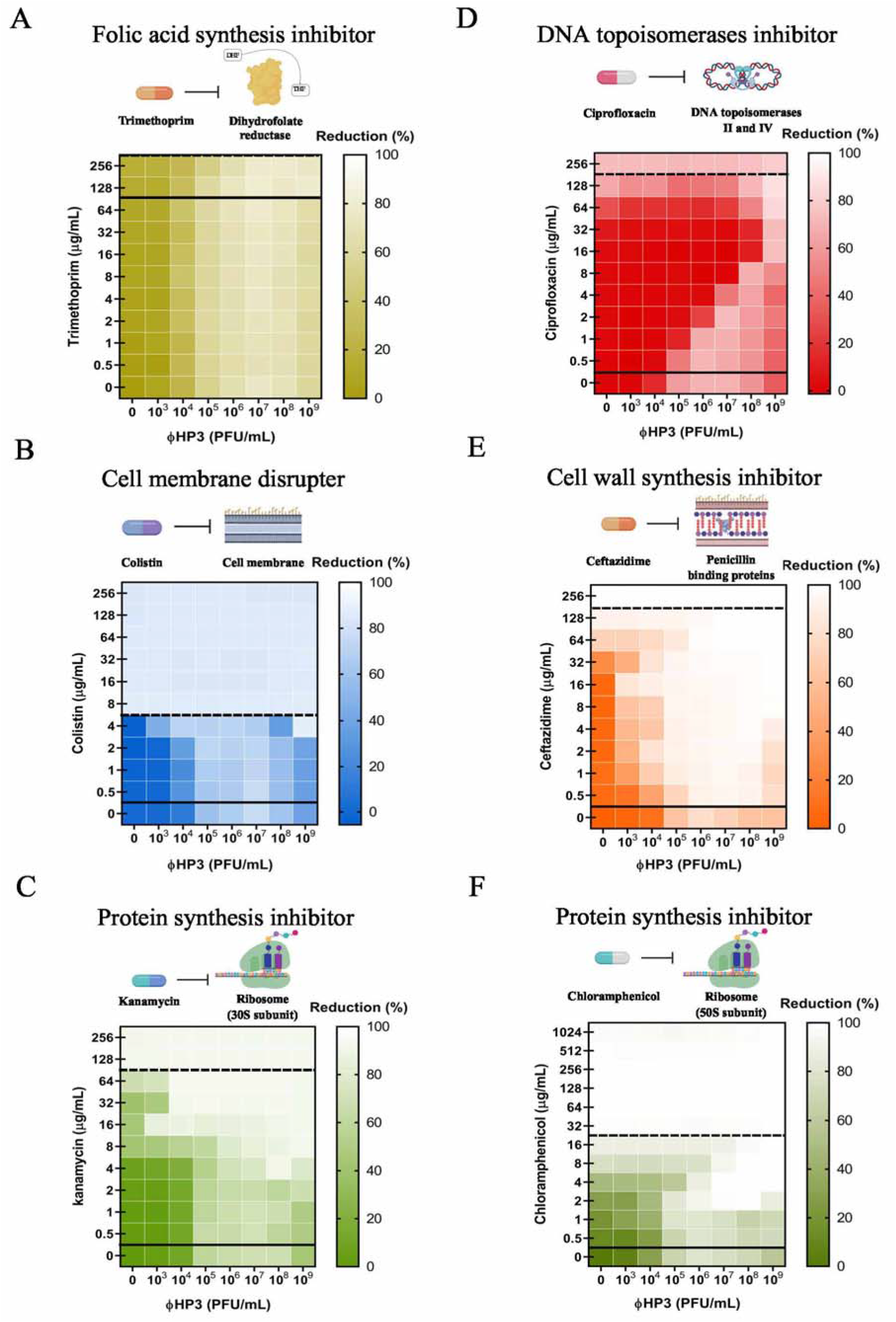
The effect of antibiotic class on phage-antibiotic synergy. A 100-fold diluted subculture of wild type JJ2528 was incubated for 4 hours, centrifuged, washed, adjusted for O.D._600nm_ of 1, inoculated onto a 96-well plate coated with HP3 and antibiotics, and the OD_600nm_ was measured every 15 minutes for a total of 24 hours with shaking. Effect of different antibiotics was studied with: (**A**) trimethoprim, (**B**) colistin, (**C**) kanamycin, (**D**) ciprofloxacin, (**E**) ceftazidime, and (**F**) chloramphenicol. Synograms represent the mean reduction percentage of each treatment from three biological replicates: *Reduction* % = [(*OD_growth control_*) – (*OD_treatment_*)]*x*100.

The cell membrane disrupter, colistin, was very effective against wild type JJ2528 in that even ≥ 8 μg/mL of colistin was able to lyse the cells almost completely (Fig. 2B). The combination of ΦHP3 and colistin produced mostly phage-dominated killing, where synergistic and antagonistic effects were only seen with one dose of 4 μg/mL and one phage titer of 10^4^ PFU/mL (Figs. 2B and 3C). In this case, unlike the cell wall biosynthesis inhibitor ceftazidime (synergism with phage) and the folic acid synthesis inhibitor trimethoprim (no effect), it seems the membrane disrupter colistin demonstrates both synergistic and antagonistic effects, similar to the chloramphenicol synograms (protein synthesis inhibitor).

**Figure 3.**
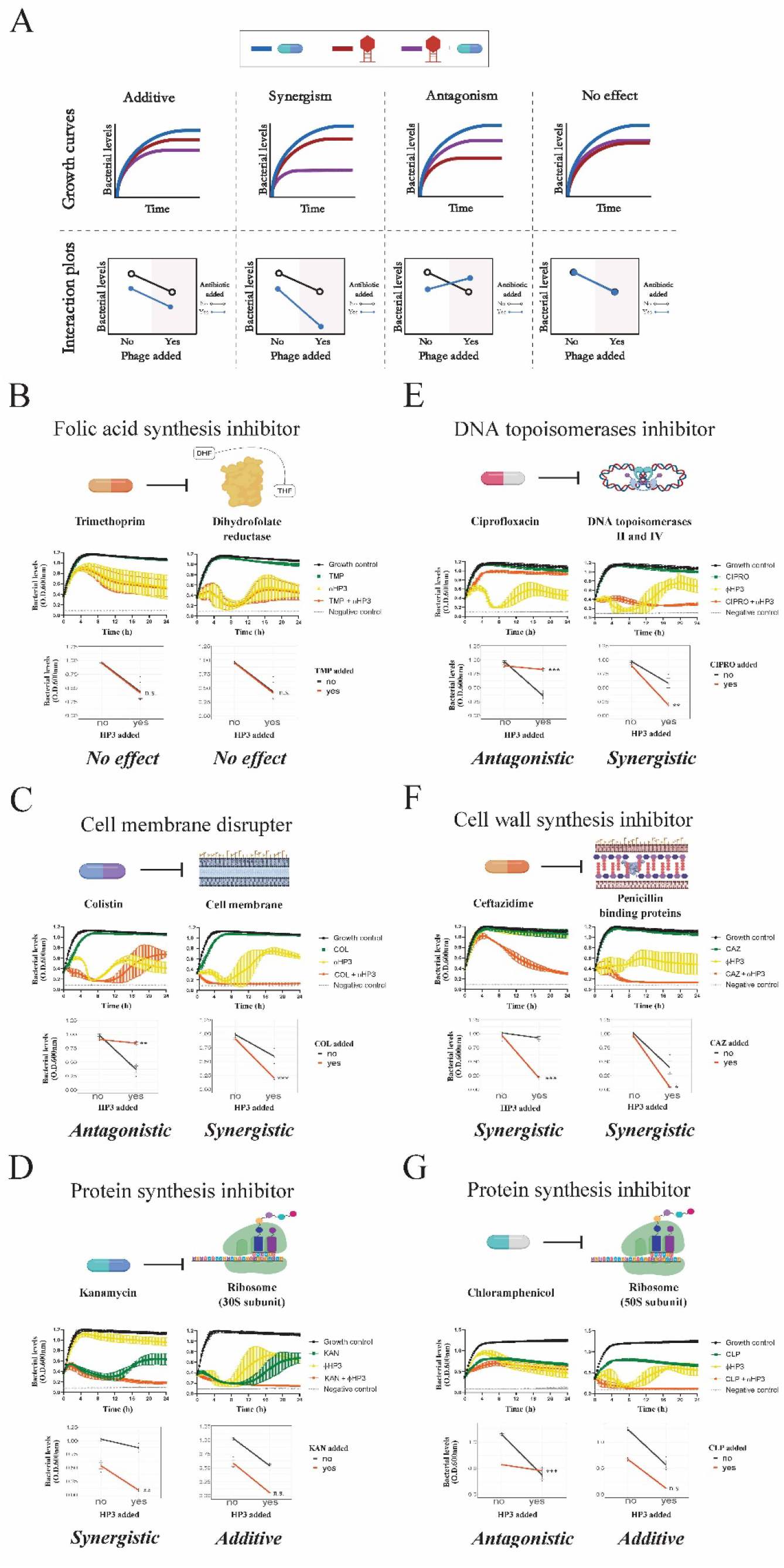
Growth characteristics and interaction plots for phage-antibiotic synergy. PAS was evaluated as described in the legends of Figure 1 and 2. Bacterial growth over time is assessed for 24 hours in the presence or absence of phage and antibiotic (top panels) and synergy assessed via interaction plots (bottom panels). (**A**) Combination of phage and antibiotic results in additive, synergism, antagonism, and/or no effect. Representative interactions between HP3 and antibiotics on wild type JJ2528 are depicted here (antibiotic dose + phage titer): (**B**) <u>trimethoprim</u>, 0.5 μg/mL + 10^5^ PFU/mL and 64 μg/mL + 10^9^ PFU/mL; (**C**) <u>colistin</u>, 4 μg/mL + 10^8^ PFU/mL and 4 μg/mL + 10^9^ PFU/mL, (**D**) <u>kanamycin</u>, 16 μg/mL + 10^4^ PFU/mL and 32 μg/mL + 10^9^ PFU/mL, (**E**) <u>ciprofloxacin</u>, 16 μg/mL + 10^8^ PFU/mL and 16 μg/mL + 10^9^ PFU/mL, (**F**) <u>ceftazidime</u>, 2 μg/mL + 10^4^ PFU/mL and 16 μg/mL + 10^9^ PFU/mL, and (**G**) <u>chloramphenicol</u>, 4 μg/mL + 10^5^ PFU/mL and 4 μg/mL + 10^9^ PFU/mL. Two-way ANOVA was employed for statistical significance testing. * P < 0.05, ** P<0.01, *** P<0.001, n.s. not significant. Growth curves show mean ± SD.

Finally, we also assessed the potential of ciprofloxacin, a DNA topoisomerase inhibitor, to act in combination with phage in killing JJ2528. Unexpectedly, the use of ΦHP3 and ciprofloxacin resulted in a highly patterned synogram (Fig. 2D) that resulted in a reduction of bacterial killing when the two agents were paired together (note the stepwise inhibition of bacterial killing as phage titers are increased). For instance, 8 μg/mL of ciprofloxacin combined with 10^7^ PFU/mL of ΦHP3 resulted in only 1% reduction, but the phage-alone treatment was around 69% reduction. This inhibition effect was then remediated by increasing the phage titer 10-fold (10^8^ PFU/mL), which led to ~69% reduction. However, this seemingly effective combination was again reversed when the ciprofloxacin doubles to 16 μg/mL, which in combination with 10^8^ PFU/mL of ΦHP3, led to only 10% reduction (Fig. 2D). Thus, the use of ciprofloxacin resulted in two outcomes, antagonism and synergism (Fig. 3E), and a pattern that was not observed for any other antibiotic class to this point. This effect is also readily apparent when examining the AUC in a violin plot (Fig. S1B, red) when compared to the other antibiotics.

### Assessment of two different antibiotics within the same mechanistic class

We next determined how the synograms may change between two different drugs that act on the same cellular pathway, in this case, protein inhibition, but have subtle mechanistic differences in their action. For this, we chose to compare the 30S ribosomal subunit inhibitor kanamycin to the 50S ribosomal subunit inhibitor chloramphenicol. Interestingly, the synogram of kanamycin treatment closely resembled the synogram produced by JJ2528 CTX-M-14 A77V/D240G treated with ceftazidime and ΦHP3 (Fig. 1F and Fig. 2C). The kanamycin synogram also demonstrated increased combinatorial efficacy that resulted in effective killing compared to either treatment alone (Fig. 2C). For instance, more than 90% reduction was observed in the combination treatment even with 4 μg/mL of kanamycin, whereas this degree of reduction in the kanamycin-alone treated cells was only found at high doses (≥ 128 μg/mL). Moreover, JJ2528 WT seems to have a subtle degree of resistance against kanamycin just as JJ2528 CTX-M-14 A77V/D240G against ceftazidime. Through growth curves and interaction plots, kanamycin and ΦHP3 were determined to act synergistically with each other in some combinations, but additive at others (Fig. 3D). Note that chloramphenicol only produced additive effects with some subtle antagonistic interactions and that its synogram was very different than that observed for kanamycin.

### Assessment of combinatorial treatment on preventing resistance

A phenomenon that was seen across some synograms is the revival of bacteria at 8 hours when high titer of HP3 (10^9^ PFU/mL) was applied as single treatment (Fig. 4A) or as combination with ineffective low dosage (0.5 μg/mL) of antibiotics (Fig. 4B). Interestingly, these “resistors”, if isolated and retested for sensitivity to phage ΦHP3, are completely recalcitrant to a second HP3 challenge, which indicates they are true resistors (data not shown). The revival was prevented when ΦHP3 was applied along with an intermediate dose (8 μg/mL) of most antibiotics, except for trimethoprim and ciprofloxacin (Fig. 4C). JJ2528 treated with high concentration (256 μg/mL) of trimethoprim and ΦHP3 still showed a second peak of growth, while high concentration of the rest of antibiotics combined with HP3 prevented this revival (Fig. 4D). Of note, phage-alone treated cells showed more fluctuations in bacterial levels, especially at later time points, compared to the positive control, antibiotic alone, and most dual treated cells. This fluctuation is more evident at higher phage titers (both phage alone and combined) seen in multiple synograms.

**Figure 4.**
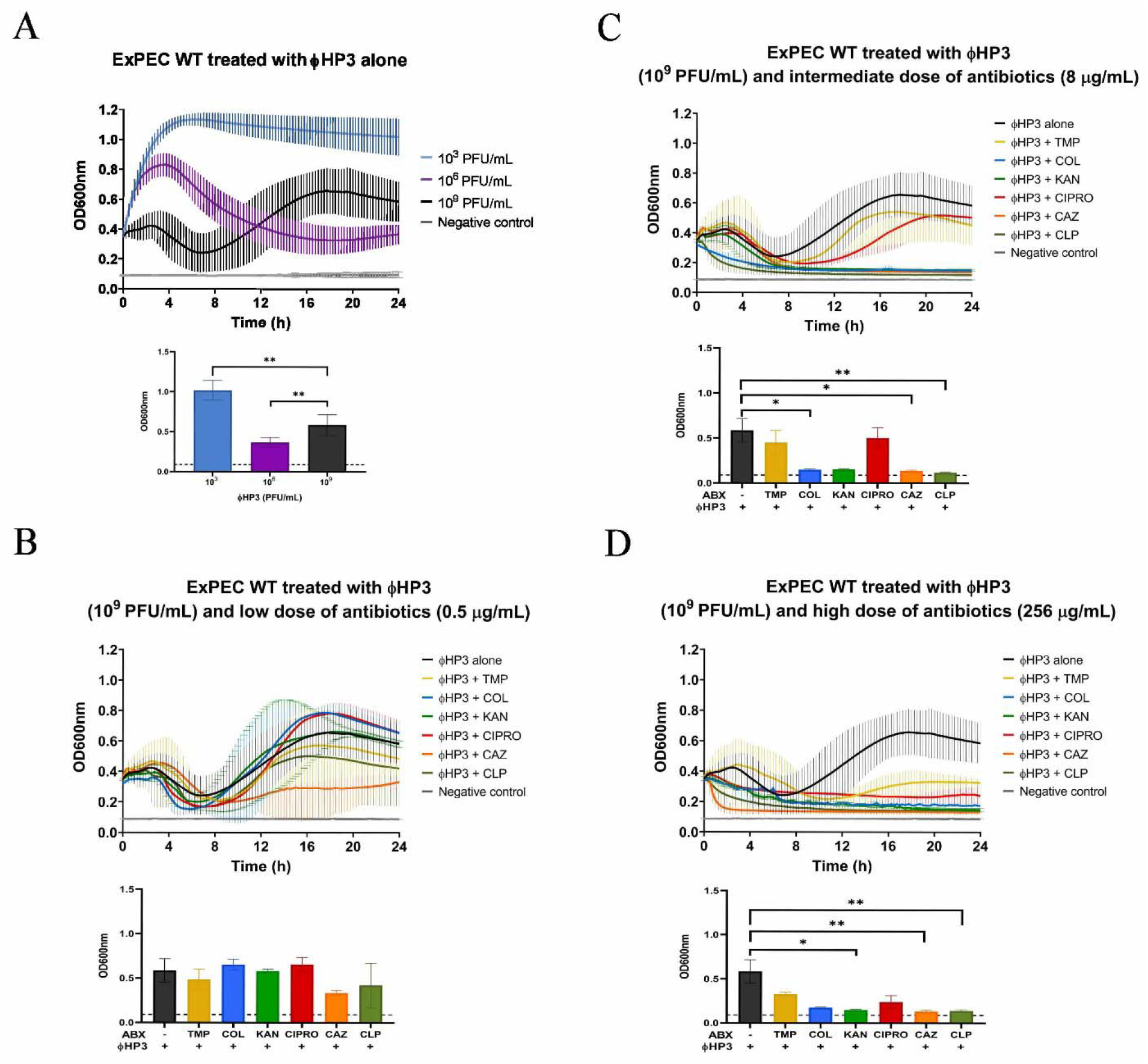
Effect of combinatorial treatment on preventing the rise of resistance. ExPEC strain JJ2528 wild type cells were treated with different titers of (**A**) ΦHP3-alone. Combination of a high titer of ΦHP3 (1×10^9^ PFU/mL) with (**B**) low, (**C**) intermediate, and (**D**) high doses of antibiotics are shown. Growth curves and bar graphs (endpoint) show mean ± SD. Kruskal-Wallis test was performed, followed by Dunn’s test for multiple comparisons. * P < 0.05, ** P<0.01, *** P<0.001, **** P<0.0001.

### Assessment of host-like environments on phage-antibiotic efficacy

The efficacy of any given antibiotic is not only influenced by the bacterium’s ability to inactivate or otherwise avoid its inhibitory effects, it also is affected by the pharmacokinetics of antibiotics in a living organism. These include the antibiotic half-life in serum or tissues, whether the host modifies or inactivates the antibiotic, as well as its oral absorption or systemic dissemination when administered. These parameters are also expected to influence phage-antibiotic synergy, as well as additional constraints the host places on phage, including, but not limited to, antagonism by the innate or adaptive immune response. To address the effect the host may have on PAS, we performed synography with two host physiologic environments important in ExPEC pathogenesis: blood and urine. In this context, blood is designed to simulate the behavior of bacteria and PAS under conditions that resemble systemic bacteremia whereas urine is meant to simulate infections of the bladder. We used human pooled urine (from multiple donors to reduce variability) and human heat-inactivated serum (to eliminate any compromising negative effects on bacterial survival caused by complement). Note that even with additional heat-inactivation, untreated bacterial levels decreased over time, especially >8h, hence we analyzed the serum synogram using the 8^th^ hour as the endpoint. We first chose an antibiotic that synergizes well with ΦHP3 against wild type JJ2528 in LB, which was the β-lactam ceftazidime (Fig. 5A). Interestingly, when this experiment was repeated with urine, the synogram was markedly different (Fig. 5B). The effective MIC of ceftazidime-alone treated cells was raised to >256 μg/mL as compared to LB. Despite being a completely different host environment with different chemical composition, a similar effect was observed when the experiment was repeated with serum (Fig. 5D). Furthermore, there was an overall less reduction (maximum 33% in urine and 26% in serum) as compared to LB (maximum 99%). When the sensitivity was increased by reducing the killing range to 50%, interactions were found in the upper right corner of each synogram with high doses of each antimicrobial (for example, synergistic in urine and additive in serum). For the serum synogram, there was consistently antagonism in the entire range of interacting regions that did not display a pattern like the ciprofloxacin synogram in LB (Fig. 5D). Since untreated bacterial levels were less in urine and serum than in the nutrient rich LB, we hypothesized that the overall reduction of killing in urine and serum is due to a lower growth rate of bacteria. Synography was performed again in both pooled human urine and human serum, but this time LB was added to the urine and serum (final concentration: 10%). The pooled human urine + 10% LB showed a maximum reduction of 70% as compared to 33% in urine-alone while the human serum + 10% LB showed a maximum reduction 28% compared to 26% in serum alone. Even though the maximum bacterial reduction in serum + LB was not significantly enhanced as compared to the urine synograms, adding LB to both urine and serum allowed synergism to appear at lower doses of dual treated cells. Moreover, by increasing the sensitivity of the synogram to a maximum 50% reduction, the urine + 10% LB synogram appears similar to the one treated in LB alone (Fig. 5A and Fig. 5C). Similarly, the serum + 10% LB synogram seems to have this trend; the antagonistic interactions in the combined treated cells appeared to have shifted toward the left side of the synogram as opposed to the serum-alone synogram (Fig. 5D and Fig. 5E).

**Figure 5.**
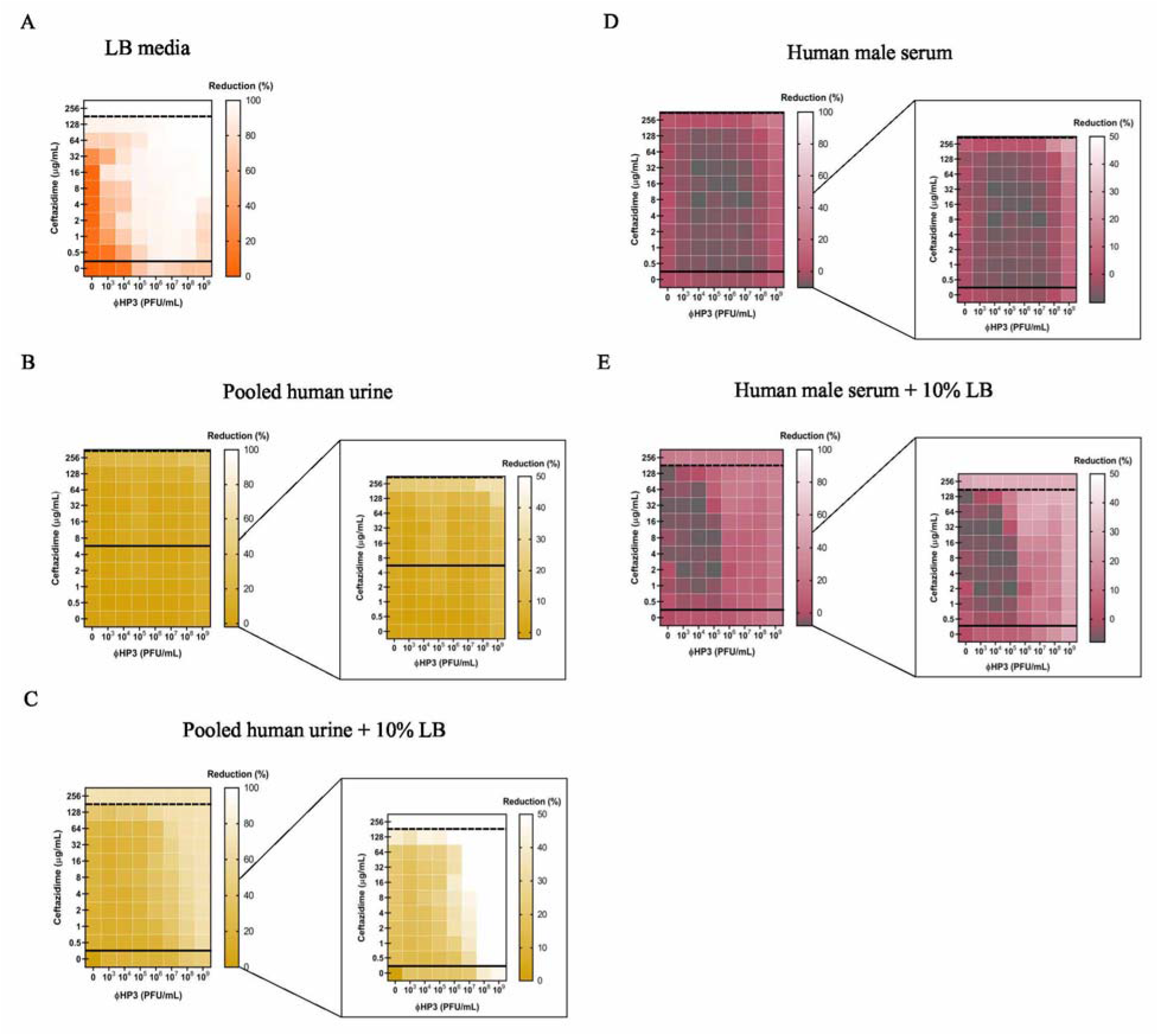
The effect of human urine and serum on phage-antibiotic synergy. A 100-fold diluted sub-culture of wild type JJ2528 was incubated for 4 hours, centrifuged, washed, adjusted for O.D._600nm_ of 1, and inoculated onto a 96-well plate coated with ΦHP3 and CAZ. The OD_600nm_ was measured every 15 minutes for a total of 24 hours for urine and 8 hours for serum with shaking. Bacterial cells are cultured in (**A**) LB, (**B**) pooled human urine, (**C**) pooled human urine + 10% LB, (**D**) human serum, and (**E**) human serum + 10% LB. Synograms represent the mean reduction percentage for each treatment in urine (N=3) and serum (N=2): *Reduction* % = [(*OD_growth control_*) – (*OD_treatment_*)]*x*100.

### Assessment of the dependence of phage type on PAS

To determine if the types of PAS observed to this point are affected by the choice of phage, we chose to perform synography with phage that is 98% identical to HP3 (termed ΦES12) but harbors a six-fold reduced burst size (60 PFU/cell for ΦHP3 compared to 10 PFU/cell for ΦES12). We examined synography when ΦES12 was combined with ceftazidime (synergism previously observed with ΦHP3) and ciprofloxacin (antagonism previously observed with ΦHP3). Unexpectedly, the combination of ΦES12 and ceftazidime resulted in a synogram with mostly additive effects and a few synergistic combinations with the overall patterning very different in composition (compare Fig. 6A with 6B). The combination of ΦES12 and ciprofloxacin yielded a synogram with some antagonistic interactions, but also yielded a quite distinct pattern compared to that observed with HP3 (Figs. 6C and 6D). In some respects, this latter synogram was more similar to the synogram observed when HP3 was combined with ceftazidime and serum (Fig. 5D).

**Figure 6.**
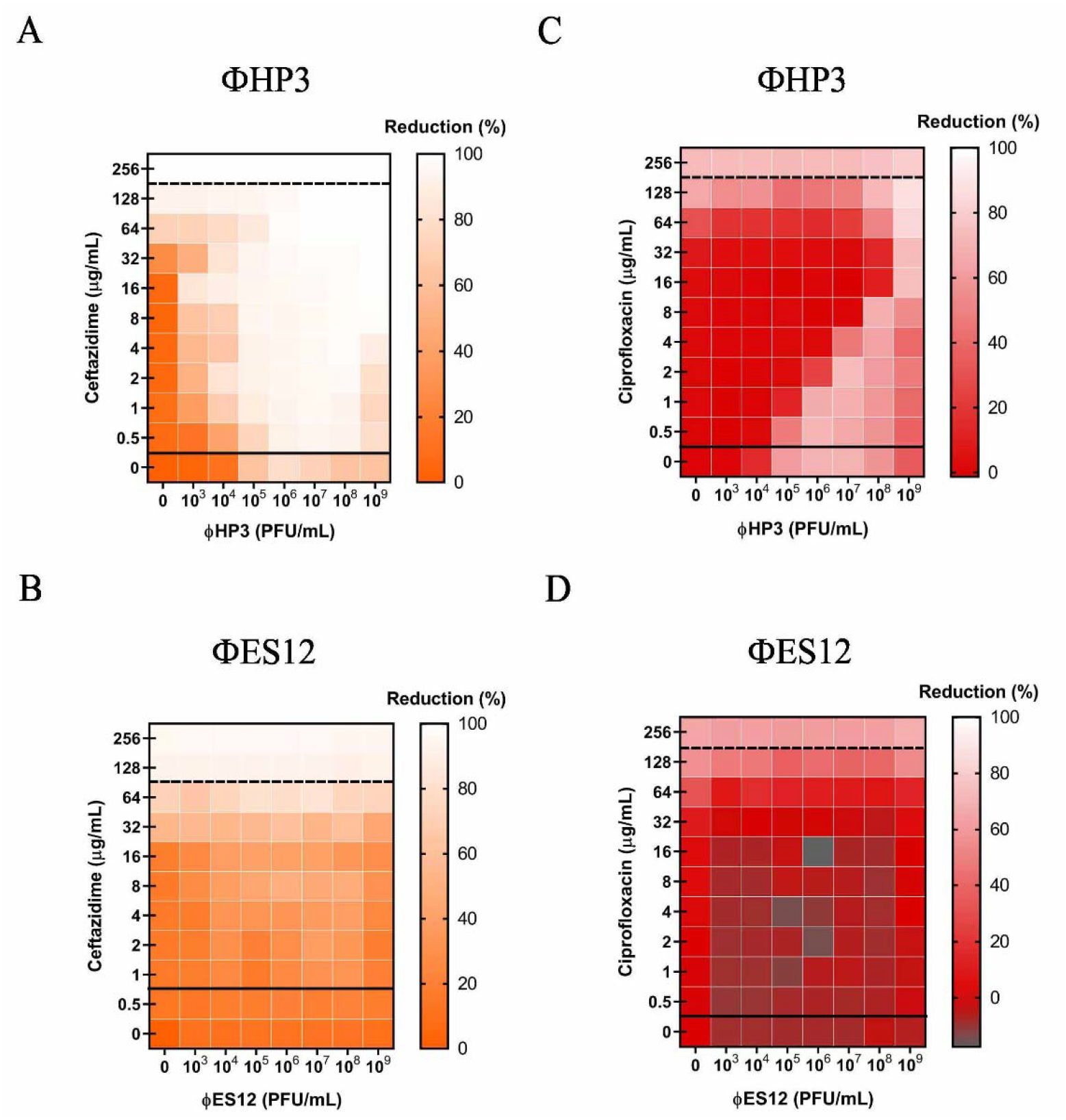
The effect of genetically similar phages on phage-antibiotic combined therapy. A 100-fold diluted sub-culture of wild type JJ2528 was incubated for 4 hours, centrifuged, washed, adjusted for O.D._600nm_ of 1, and inoculated onto a 96-well plate coated with phages (ΦHP3 and ΦES12) and antibiotics in LB media. OD_600nm_ was measured every 15 minutes for a total of 24 hours with shaking in between. Synograms show wild type ExPEC treated with (**A**) ΦHP3, and (**B**) ΦES12 with antibiotics (ceftazidime and ciprofloxacin). Synograms represent the average reduction percentage for each treatment from three biological replicates: *Reduction* % = [(*OD_growth control_*) – (*OD_treatment_*)]*x*100

## Discussion

Recognizing that a phage used in combination with antibiotics might yield possible beneficial interactions that can enhance in vivo efficacy of the antimicrobials, improve clinical outcomes, and decrease resistance, we tested a recently discussed phage highly specific against the pandemic ST131 clonal group of *E. coli* for synergistic interactions with all major classes of antibiotics. Primary findings from our study reveal: (i) the development of a new, high-throughput platform that quickly assesses the effect of various phage and antibiotic concentration on bacterial growth, an analysis we call synography (and the resulting data represented in a synogram); (ii) that synograms demonstrate a wide range of conditions in which combinatorial treatment is synergistic, additive, or antagonistic, sometimes all three present in the same analysis; (iii) that a phage may demonstrate highly effective killing or inhibition when combined with one class of antibiotics but may lack this same effect when combined with another class; (iv) that phage may restore the competency of antibiotics even in bacteria that encode resistance elements against the chosen antibiotic, an effect we term “phage adjuvation” because the phage adjuvates, or make better, the antibiotic; (v) that highly genetically similar phages produce dramatically different synograms even when these are combined with the same class of antibiotics; (vi) that phage-antibiotic synergy may prevent resistance, but only when the antibiotic concentration is increased; (vii) and finally, the host-like conditions substantially influence PAS, and the synogram profiles in general, thus reflecting the need to test such antibacterial effects under conditions that more reliably simulate the host environment. In this case, the dampening of PAS seems to be due to a reduced growth rate when the bacterium is in urine or blood. Collectively, our work suggests that real-time assessment and screening of unique phage-antibiotic combinations (and therapeutic doses) is possible and will likely substantially influence both the choice of the antibiotic and the phage, as well as their doses, to improve clinical outcomes.

### Efficacy of combinatorial treatment in wild type and antibiotic-resistant bacteria

An area of phage therapy that is in its infancy is whether phage can help re-sensitize resistant bacteria toward antibiotics. Often, the options for antibiotics can be limited by bacterial resistance or by the patient’s own medical conditions. In this investigation, we studied the efficacy of combinatorial therapy using antibiotics that can be targeted by bacterial enzymes; these antibiotics are rendered ineffective if they were to be used alone. We found that the MIC was lowered for both wild type and resistant JJ2528 when the antibiotic was combined with phage. In other words, phages seemed to act as a type of adjuvant to antibiotics, much like we consider alum for antigens in vaccines, even with bacteria harboring genetic elements that confer resistance to the antibiotic. In this sense, they “resensitize” bacteria by lowering the dose of antibiotic needed to achieve a similar reduction of bacterial levels compared to the antibiotic-alone. There are other ways in which bacteria can be “resensitized”, for instance, through transduction whereby sensitiving genes are delivered to bacteria after phage infection (29). As noted here, the environmentally isolated phage in our study is already very efficacious in resensitizing a multi-drug resistant JJ2528 without prior transduction events. In this regard, it seems that such an effect is heavily dependent on finding the “optimal” dose of both agents such that their individual molecular mechanism of action do not interfere with each other. In addition, synograms between resistant and wild type JJ2528 showed similar types of interactions in general. For example, if synergism is found between a particular combination of phage and antibiotic in wild type bacteria, just as the case of HP3 and ceftazidime, this synergism also extends to resistant bacteria. The same applies to the additive-antagonistic effect found in both chloramphenicol-sensitive JJ2528-WT and chloramphenicol-resistant JJ2528-CAT. These results imply that the stoichiometry of each agent (e.g. the relative molar amounts of each) is an important determinant of activity, and can be used to overcome the genotype of the pathogenic bacterium.

### The class of antibiotic determines the type of interaction with phage

The possible types of interactions between phage and antibiotic were largely dictated by the class of antibiotics employed during therapy. In our system of wild type JJ2528 treated with the same phage but with different antibiotics, we i) observed additivism, synergism, antagonism, and neutrality; ii) found that synergistic effects with an antibiotic does not translate to a different antibiotic of the same class that targets similar bacterial pathways; and iii) more than one type of interactions can exist simultaneously. Since there is a myriad of interactions between phage and antibiotic, there lies a possibility that the interference of some bacterial processes by certain antibiotics may also affect the lytic cycle of the phage. For instance, the synergism and antagonism seen in the membrane disrupter colistin can be either caused by the complexity of the drug (major and minor forms with amphiphilic property) or by its primary interaction with LPS that leads to cell membrane destabilization (30, 31). Since many phages use LPS as a receptor on the bacterial cell, it is possible that the synogram reflected colistin affecting the initial adsorption stage of the phage (32, 33). Another example is the synergistic interaction found between ΦHP3 and the cell wall inhibitor ceftazidime, which can be correlated to the increased phage production caused by cephalosporins. These antibiotics have been demonstrated to increase cell filamentation as well as to cause the production of more phage particles (21, 34–36). It is also possible that the combined action of phage-derived products, which can disrupt the bacterial membrane integrity, along with the action of ceftazidime on the cell wall, may cause a fragile barrier that allows easier cell lysis. The dose-dependent pattern between ciprofloxacin and HP3 is a clear antagonistic behavior that can be explained by the primary targets, DNA topoisomerases, that are involved in DNA replication encoded by both bacterium and phage (37). ΦHP3, used in this study, encodes two subunits of DNA topoisomerase II (data not shown), suggesting that ciprofloxacin inhibits both the bacterial and the phage topoisomerases. Similarly, recent biofilm PAS studies noted that sequentially treating cells with phage and ciprofloxacin (noted synergism) instead of a simultaneous application (noted antagonism), may have allowed phage replication to occur first before ciprofloxacin’s interruption (23, 38). These results raise the possibility that the type of interactions in each phage-antibiotic combination is heavily dictated by the primary target of the antibiotic and the cellular processes required for phage replication (34, 39, 40). Lastly, since protein synthesis inhibitors most likely would interfere with phage production, the dominant synergistic effects seen in kanamycin was unexpected. Other PAS studies with protein synthesis inhibitors have also found synergistic interactions *in vitro* (39, 41). Thus, we speculate that HP3 possesses a mechanism to bypass the antibiotic inhibition of the ribosome, and allow synthesis of phage proteins. This notion is supported by the discovery of phage-encoded ribosomal subunits (42). The bactericidal property of kanamycin may have contributed to the enhanced killing found only in the kanamycin synogram. Bactericidal agents, like kanamycin, accelerate cellular respiration rate, followed by stimulation of hydroxyl radicals that are thought to cause cell death (43). Since phage replication is thought to rely on metabolically active bacteria, increasing cellular respiration may enhance phage-mediated killing (44, 45).

### PAS prevents development of phage resistance

From an evolutionary standpoint, the imposition of two different selective pressures on bacteria may reduce the chances for them to develop potential resistance (46). In our study, it was observed that combinatorial treatment prevented the rise of secondary bacterial growth that was often observed in phage-only treated cells at later time points. This is consistent with studies that have demonstrated that only combinatorial therapy can effectively prevent the rise of phage-resistant variants (47–49). Such an observation bodes well for the prospects of using PAS to prevent resistance.

### The host physiological environment changes the efficacy of PAS

Susceptibility testing involves tightly regulated parameters that include media, bacterial inoculum, dilutions of antibiotic, as well as the plate used for the assay (50). However, *in vitro* testing may not always correlate to *in vivo* efficacy. An inoculum effect was discovered to be an *in vivo* concern, and this represents only one of the tightly regulated parameters (51). Generally speaking, for all antibiotics tested here (except for trimethoprim, which was ineffective), as the concentration of bacteria increases, the effectiveness of the antibiotic decreases, consistent with the antibiotic being slowly dosed out (Fig. S2). To study the potential impact that the physiological environment may have on phage-antibiotic interactions, medical simulators such as human urine and human serum were employed in place of the LB media. Urine has been shown to increase the apparent MIC of *E. coli* to several classes of antibiotics (52), and this phenomenon was observed here. We also found that urine affected the efficacy of phages. Only the dual treatment showed an effective reduction of bacterial levels and synergism was preserved, albeit modestly. This suggests that phage and ceftazidime are also effective in synergizing with each other even in acidic environments like urine. Since phages are routinely used to treat UTI at the Eliava Institute in Georgia, and there are increasing reports of successful phage therapy on UTI, more investigation is needed to understand the parameters required for successful treatment in the urine environment (53, 54). Phages face additional challenges in a complicated host system like the blood that may lead to phage inactivation. In our serum synogram, there were antagonistic interactions observed only in combined treatment that can only be overcome by higher doses of ceftazidime and higher phage titers. It appears that serum does not cause ceftazidime to antagonize phage-killing the way that ciprofloxacin does in LB. Conversely, the effect seems to be more of a failure to inactivate bacteria when both antimicrobials are present in serum, as opposed to single agents, and highly dependent on active bacterial growth. It would be of interest to screen for phages that somehow activate bacterial growth.

### Similar, but distinct, phages result in different synograms

Phages that are chosen for therapy should be carefully characterized to avoid the presence of potentially harmful genes that yield toxins, antibiotic resistance, and virulence to the bacterial host. From our recent characterized phage library (55), we chose to study antibiotic-phage interactions using ΦES12. Similar to ΦHP3, this phage is devoid of potentially harmful genes, and its genome sequence is 98% identical to ΦHP3. They have similar genome size (ΦHP3=168kb, ΦES12=166 kb), G + C content (ΦHP3=35.4%/o, ΦES12=35.37%), number of open reading frames (ΦHP3=274, ΦES12=267), number of tRNAs (ΦHP3=11, ΦES12=9), similar absorption within 10 minutes (ΦHP3=98%, ΦES12=93%), and similar latent periods (ΦHP3=22.5 min, ΦES12=26 min), but they differ in burst size (ΦHP3=60 PFU/mL, ΦES12=9.6 PFU/mL)(55). Since the burst size is high in ΦHP3 and low in ΦES12, and burst size is one of the factors that affect phage-mediated killing, we hypothesize that the differences in synogram observed between these phages might be due to this factor. This is especially true when only phage-alone treated cells are considered (Figs. 6A and 6B). In ΦHP3-treated cells, there is a high reduction of bacterial levels at t > 4h, but this is absent in ΦES12-treated cells. In the synograms, this is represented by consistently high bacterial levels, or darker color, in ΦES12-alone treated cells in contrast to ΦHP3-treated cells. Thus, it seems burst size is one important determinant in achieving synergism, with phage exhibiting a greater burst size being preferable.

Overall, our investigation of phage-antibiotic interactions paves a new path to explore the complexity of success of dual therapy, especially in physiological conditions. We developed a rapid *in vitro* assay that can be employed to assess the types of interactions between phage and antibiotic before translating this knowledge into a combined therapy. Our work demonstrates (1) how different interactions between phage and antibiotic are strongly affected by the class of antibiotics, (2) how phage generally lower the MIC of the antibiotic, (3) how phage and antibiotics suppress resistance, (4) how bacterial resistance toward antibiotic impacts the combination therapy, (5) how host factors like urine and serum affect these types of interactions, and (6) how similar phages may result in dramatically different outcomes. Future work should lead to understanding how the efficacy of a phage cocktail combined with antibiotics alters efficacy, especially in complex host systems. In addition, determining the effect of simultaneous vs sequential treatment to reduce antagonistic interactions, as well as the actual mechanisms behind each synergistic and antagonistic effect in combinatorial treatment, are fertile grounds for future research.

## Materials and methods

### Bacterial culture, plasmids, phage and antibiotics

The clinical isolate ExPEC ST131 strain JJ2528 used in this study was kindly provided by Dr. James R. Johnson (56) and the plasmids (pTP123-CTX-M-14 WT and pTP123-CTX-M-14 A77V/D240G) that confer antibiotic resistance were kindly provided by Dr. Timothy Palzkill (25). The phages used in this study, ΦHP3 and ΦES12, were previously isolated from environmental sources (24, 55). All antibiotics were prepared fresh, and filter sterilized (except chloramphenicol and trimethoprim due to solvent used for these antibiotics). Information about maintenance of bacterial culture, phage purification, classes of antibiotics and solvent employed for antibiotics can be found in the Supplementary Methods.

### Synergy testing in LB media, human urine, and human serum

Synergy testing was performed with LB media, human pooled urine, and commercial human serum. A subculture of *E. coli* strain JJ2528 was incubated for 4 hours in LB, centrifuged, washed, and recentrifuged. The pellet was resuspended in the medium under test, adjusted to ~ 1 x 10^9^ CFU/mL (OD_600nm_ = 1), and 100 μL was inoculated into each well of the micro-titer plate that contained the checkerboard of phage and antibiotic concentrations (50 μL for each antimicrobial). The OD_600nm_ was measured every 15 minutes at 37°C for a total of 24 hours with continuous shaking in a Biotek Synergy HT (Biotek, VT, USA). For more details on synergy assays, urine collection, and serum information, visit Supplementary Methods.

### Inoculum effect

To determine the effect of inoculum size on the efficacy of antibiotics, 2-fold serial dilutions of antibiotic were added to a 96-well plate. A bacterial sub-culture of JJ2528 was prepared as described for synergy testing, and serial 10-fold dilutions were inoculated into the microtiter plate, and grown for 24 hours at 37°C in a shaker. For each bacterial inoculum, the well with the lowest antibiotic concentration that showed bacterial clearance was marked as the MIC (minimum inhibitory concentration). For more details on antibiotics and bacterial dilution, visit Supplementary Methods.

### Data representation and statistical analysis

To generate synograms, absorbance readings from three biological replicates were normalized with the negative control, and the treated wells were deducted from the positive control (no treatment) to yield percent reduction.

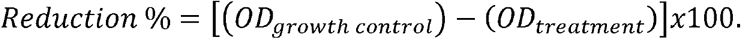

Two-way ANOVA was employed on interaction plots to analyze possible synergism between phage and antibiotics. AUC was generated for the interacting region of each synogram and normalized. Figures were generated with Biorender. For more details on the statistics, see Supplementary Methods.

## Acknowledgments

We thank Dr. James R. Johnson for the ExPEC clinical isolate JJ2528 and Dr. Timothy Palzkill for the beta lactamases pTP123-CTX-M-14 WT and pTP123-CTX-M-14 A77V/D240G used in this study.

