## Supplementary Information for "Phage-Antibiotic Synergy is Driven by a Unique Combination of Antibacterial Mechanism of Action and Stoichiometry"

**Supporting Information Appendix**

**Supplementary materials and methods**

### *Bacterial strain, culture, and plasmids*

All strains were stored in LB medium with 25% (vol/vol) glycerol at -80°C. Periodically, a bacterial plate stock is made by streaking glycerol stock onto LB-Miller agar plates (Fisher Bioreagents, ON, Canada) for isolation of single colonies. Several colonies from a bacterial plate stock were used to inoculate 5 mL of LB for an overnight culture. For 10-fold serial dilution plating of bacterial cultures, the track dilution technique was performed on LB-Miller agar (Fisher Bioreagents, ON, Canada) (57). Lastly, pTP123-CTX-M-14 WT and pTP123-CTX-M-14 A77V/D240G were individually introduced into ExPEC JJ2528 via electroporation at 1.8kV (Eppendorf Eporator). All bacterial cultures were grown with shaking at 37°C.

### *Phage purification and storage*

Preparation of phage stocks involves phage amplification, phage precipitation with NaCl and 7.5% PEG8000, clean-up with DNase & RNAse (final 5µL/mL), phage purification with CsCl gradient centrifugation (30k rpm, 10°C, 18 hours) and dialysis in phage buffer. Purified phage stocks were stored in a 5 mL glass vial at 4°C. The titer of the phage stock was determined by spotting 5 µL of each dilution (10-fold) onto an LB-Miller plate that was already coated with a top agar layer (7.5%) mixed with 100 µL of overnight bacterial culture.

### *Antibiotics and solvent*

The following antibiotics and solvents were used in this study: chloramphenicol (Sigma Aldrich, MO, USA) dissolved in ethyl alcohol (final <1%), ceftazidime hydrate (Sigma Aldrich, MO, USA) dissolved in saturated NaHCO_3_ solution (final <1%), ciprofloxacin hydrochloride (Corning, VA, USA) dissolved in ddH_2_O, kanamycin sulfate (Omniflur) dissolved in ddH_2_O, colistin sulfate salt (Sigma Aldrich, MO, USA) dissolved in ddH_2_O, and trimethoprim (Sigma Aldrich, MO, USA) dissolved in DMSO (final <1%).

### *Synergy testing in LB media*

To start a sub-culture, 100 µL of the overnight culture was inoculated into 10 mL of LB. At the end of 4 hours of incubation, this sub-culture was centrifuged (4.4k rpm, 24°C, 10 min.), washed, and centrifuged again (4.4k rpm, 24°C, 10 min.). The supernatant was discarded, and the bacterial pellet was resuspended with LB media. Later, 100 µL of the bacterial culture (OD_600nm_ = 1) was inoculated into each well of the 96-well-plate (Genesee Scientific, CA, USA) that was previously coated with varying concentrations of phage (50 µL, final 10^3^-10^9^ PFU/mL) and antibiotic (50 µL, final 0.5-256 µg/mL for most antibiotics and 0.5-1024 µg/mL for chloramphenicol) for a final 5 x 10^8^ CFU/mL of bacteria per well. The multiplicity of infection (MOI) ranges from 2 x 10^-6^ to 2. Synergy testing was performed with a Biotek Synergy HT (Biotek, VT, USA), in which the OD_600nm_ was measured every 15 minutes for a total of 24 hours with shaking in between.

*Urine collection and heat-inactivation of serum*

Human urine was collected and pooled from both male and female donors. Donors refrained from consuming caffeine or medications the day prior to collection, and potential hormonal effects were minimized. Human male serum (Type AB, heat-inactivated, off-the-clot serum, Access Biologicals LLC, California, USA) was further heat-inactivated for an additional hour and employed for synergy testing.

### *Pooled human urine synergy test*

Urine from female and male donors were warmed to 37°C, filtered, and stored at -20°C. To thaw urine prior to synergy testing, samples were placed in a 37°C incubator and allowed to thaw for 4-6 hours with agitation every hour. Then, urine from three male donors and two female donors were pooled into one single sample (pH ~5.73). Synergy testing with pooled human urine was performed just as previously described. Briefly, sub-culture was grown in LB for 4 hours, centrifuged, washed with urine, resuspended in urine, and adjusted for O.D_600_ nm = 1. This culture was inoculated into a 96-well plate that was previously coated with antibiotic and phage. The OD_600nm_ was measured every 15 minutes for a total of 24 hours with continuous shaking. Three biological replicates were performed. For synergy testing on human pooled urine + 10% LB, assay was carried out as mentioned but media had a final 10% LB per well.

### *Human heat-inactivated serum synergy test*

Human serum was stored at -20°C and aliquots were thawed overnight at 4°C. Prior to synergy testing, human serum was further heat-inactivated for an additional hour at 56°C. Synergy testing with human serum was performed as previously described. Briefly, sub-culture was grown in LB for 4 hours, centrifuged, washed with PBS, resuspended in serum, and adjusted for O.D_600_ nm = 1. This culture was inoculated into a 96-well plate that was previously coated with antibiotic and phage. The OD_600nm_ was measured every 15 minutes for a total of 8 hours with shaking in between since bacterial growth in the positive control seems to destabilize and decrease over time even with further heat inactivation (> 8h). Two biological replicates were performed. For synergy testing on human serum + 10% LB, assay was carried out as mentioned but media had a final 10% LB per well.

### *Inoculum effect*

To test the effect of the inoculum on the efficacy of the antibiotics, 100 µL of the overnight culture was inoculated into 10 mL of LB and cultured for 4 hours. Then, the sub-culture is centrifuged, washed, and resuspended with LB as described. The bacterial culture was then adjusted to O.D._600nm_ = 1.2 and diluted 10-fold. Diluted bacteria (50 µL) was added to a 96 well plate (Genesee Scientific, CA, USA) that was previously coated with antibiotic (50 µL, final 1-512 µg/mL) for a final bacterial concentration of 10^3^-10^9^ CFU/mL.

### *Data representation and statistical analysis*

First, data was normalized by deducting the absorbance of the negative control from each endpoint of the positive control, phage only, antibiotic only, and phage-antibiotic treated cells. Then, cells that underwent treatment (both single and combined) were deducted from the positive control to yield the percent reduction, which is represented in visual format with a heatmap.

To analyze possible synergism between phage and antibiotic, both interaction plots and two-way ANOVA were employed for the endpoints of each treatment. Interaction plots showed whether two individual agents interact with each other, and the two-way ANOVA test was performed to see whether the visual representations were statistically significant (58). To account for the reduction in bacterial levels over time, area under the curve (AUC) was computed for the regions of the synograms and AUCs were normalized according to the lowest and highest mean of three biological replicates.

**Supplementary Figures**

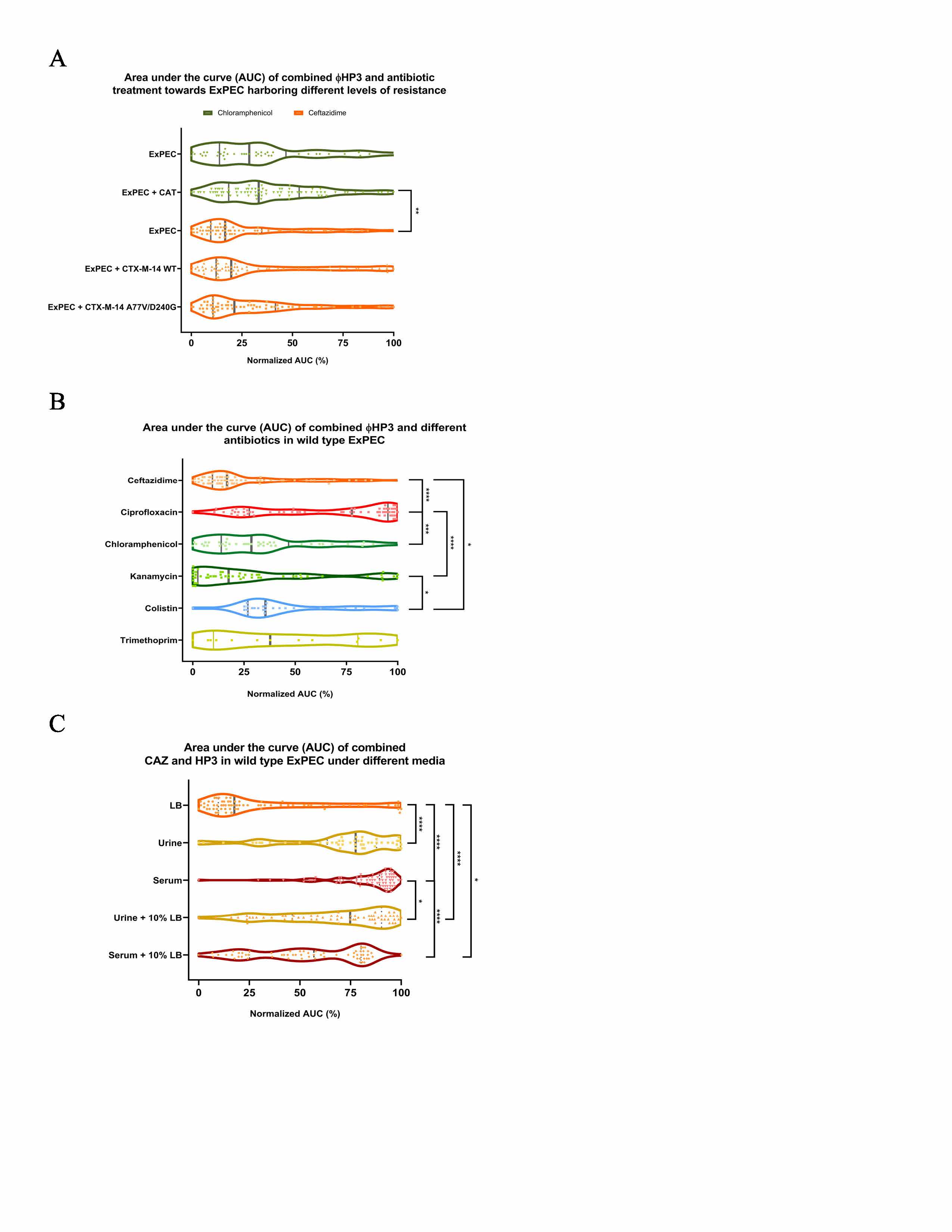

**Supplementary figure 1. Violin plots for the interacting region of the synograms.** Area under the curve (AUC) for the possible interacting region of the synograms are shown in violin plots where the median is the solid thick vertical line and the quartiles are solid thin lines. **(A)** Comparison between bacterial resistance, **(B)** comparison between classes of antibiotics, and **(C)** comparison between media. AUCs were normalized. Kruskal-Wallis test was performed, followed by Dunn’s test for multiple comparisons. * P < 0.05, ** P<0.01, *** P<0.001, **** P<0.0001

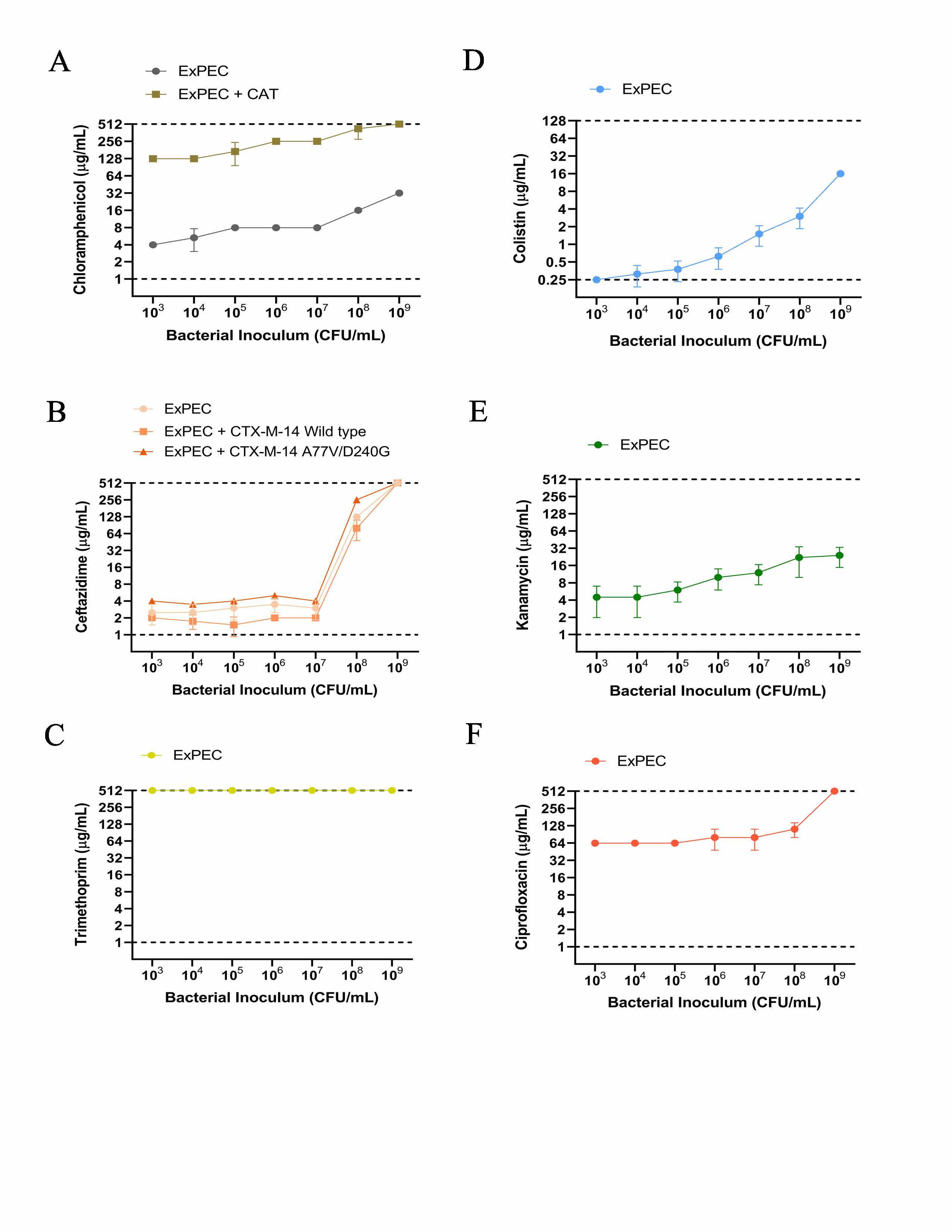

**Supplementary figure 2. Inoculum effect of different antibiotics.** A 100-fold diluted sub-culture of ExPEC (strain JJ2528) was incubated for 4 hours, centrifuged, washed, adjusted for O.D._600nm_ of 1.2, serially diluted, and inoculated onto a 96-well plate coated with different antibiotic concentrations. The lowest dose of antibiotic that inhibited visible bacterial growth was marked as the MIC for each bacterial inoculum. Inoculum effect was tested against different bacterial transformants with ceftazidime and chloramphenicol: **(A)** chloramphenicol on JJ2528-WT and JJ2528-CAT, and **(B)** ceftazidime on wild type JJ2528, JJ2528 CTX-M-14 wild type, and JJ2528 CTX-M-14 A77V/D240G. Inoculum effect was also tested on JJ2528 wild type with different antibiotics: **(C)** trimethoprim, **(D)** colistin, **(E)** kanamycin, and **(F)** ciprofloxacin. Dashed lines represent the limit of detection.

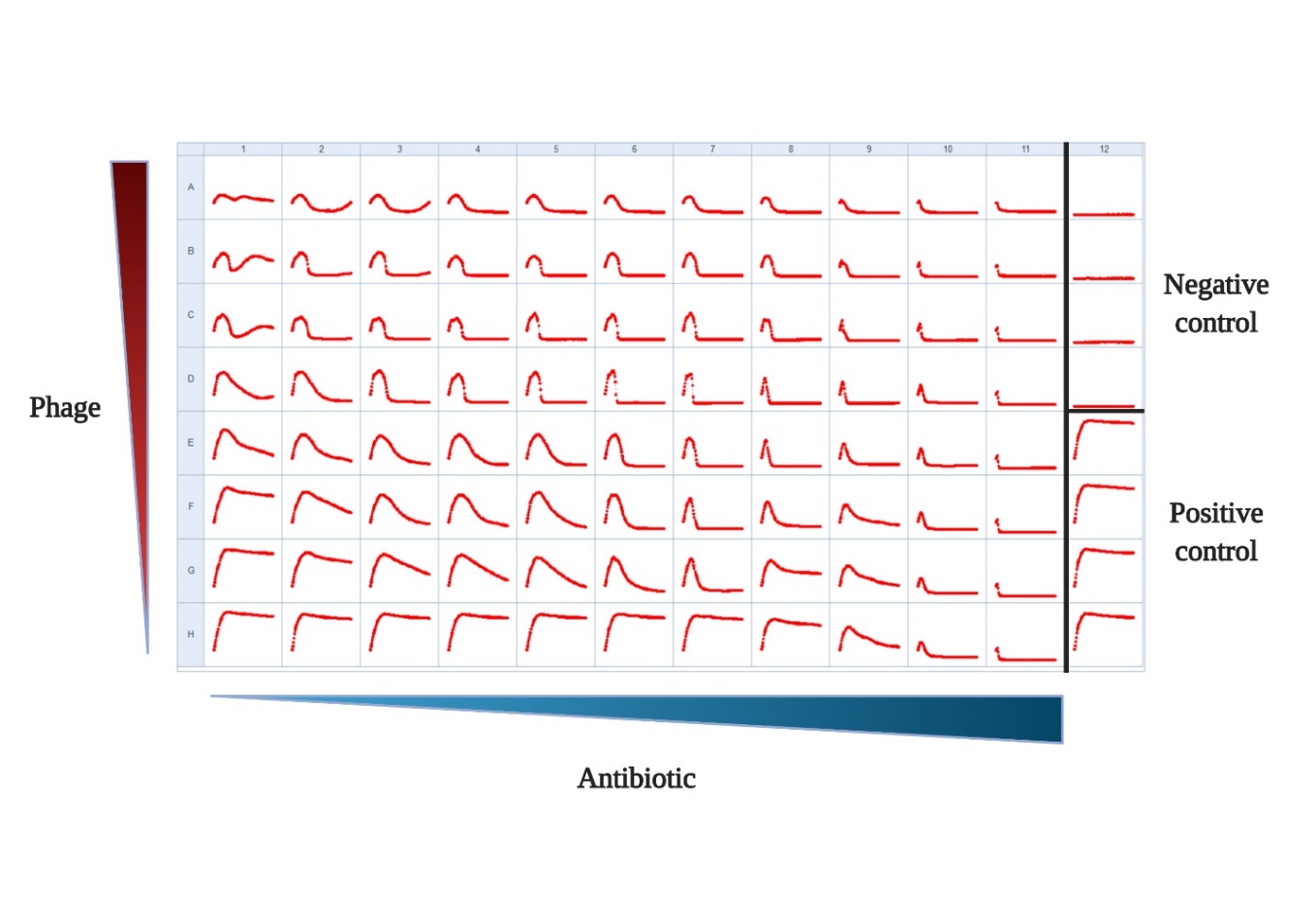

**Supplementary figure 3. Diagram of a 96-well plate showing raw absorbance readings for the combinatorial treatment on wild type JJ2528.** A 100-fold diluted sub-culture of ExPEC (strain JJ2528) was incubated for 4 hours, centrifuged, washed, adjusted for O.D._600nm_ of 1.0, and inoculated onto a 96-well plate coated with different antibiotic concentrations and phage titers. The plate shows the raw absorbance readings for one biological replicate.
